# 14-3-3γ prevents centrosome duplication by inhibiting NPM1 function

**DOI:** 10.1101/2019.12.19.883264

**Authors:** Arunabha Bose, Kruti Modi, Suchismita Dey, Somavally Dalvi, Prafful Nadkarni, Mukund Sudarshan, Tapas Kumar Kundu, Prasanna Venkatraman, Sorab N. Dalal

## Abstract

14-3-3 proteins bind to ligands via phospho-Serine containing consensus motifs. However, the molecular mechanisms underlying complex formation and dissociation between 14-3-3 proteins and their ligands remain unclear. We identified two conserved acidic residues in the 14-3-3 peptide-binding pocket (D129 and E136) that potentially regulate complex formation and dissociation. Altering these residues to Alanine led to opposing effects on centrosome duplication. D129A inhibited centrosome duplication while E136A stimulated centrosome amplification. These results were due to the differing abilities of these mutant proteins to form a complex with NPM1. Inhibiting complex formation between NPM1 and 14-3-3γ led to an increase in centrosome duplication and overrode the ability of D129A to inhibit centrosome duplication. We identify a novel role of 14-3-3 in regulating centrosome licensing and a novel mechanism underlying the formation and dissociation of 14-3-3 ligand complexes dictated by conserved residues in the 14-3-3 family.

## Introduction

Partitioning of DNA into two daughter cells relies on the accurate organization of microtubules into a spindle (1). Spindle organization in mammalian cells is primarily dependent on the centrosome, which consists of two centrioles, a mother centriole, and a daughter centriole, surrounded by the pericentriolar matrix (PCM) (2-4). The centrosome duplicates only once during a cell cycle (5) generating two centrosomes that nucleate a bipolar mitotic spindle (1,6). Centrosome amplification is observed in human tumors (reviewed in (7,8)) and an increase in centrosome number leads to an increase in aneuploidy (8-11) and invasion (12). A loss of centrioles leading to a decrease in centrosome number is also associated with chromosomal instability and aneuploidy (13,14). Therefore, changes in centrosome number can have adverse consequences for the cell.

At the end of mitosis and at the beginning of G1, the two centrioles in a centrosome are disengaged due to the Polo-Like Kinase 1 (Plk1)- and Separase-induced degradation of the inter-centriolar linker (15-18). Disengagement is a licensing event for centrosome duplication, as centrioles do not duplicate unless they are ∼80 nm apart (16,19). Once centrioles are disengaged, CDK2 activation leads to the initiation of procentriole biogenesis (20) and CDK2 overexpression results in centrosome amplification (20,21). CDK2 regulates procentriole biogenesis in part by phosphorylating the inter-centrosomal linker protein Nucleophosmin (NPM1), on a Threonine residue at position 199 (T199) (22,23). NPM1 associates exclusively with unduplicated centrosomes and expression of a phospho-site mutant of NPM1, T199A, results in the retention of NPM1 at the centrosome and inhibits centrosome duplication (22,23). In contrast, the expression of a phospho-mimetic mutant, T199D, leads to centrosome amplification (24). During S-phase, Polo-Like Kinase 4 (Plk4) recruitment at the site of procentriole biogenesis initiates centriole duplication (25,26). Overexpression of Plk4 results in centrosome amplification and its depletion results in a loss of centrioles (27,28). During G2, pro-centrioles elongate and the mother centrioles mature and acquire distal and sub-distal appendages and the ability to nucleate PCM (5,21,26,28,29). Before entry into mitosis, CDK1 mediates the separation of centrosomes to the two poles, leading to the formation of a bipolar spindle (29-31).

14-3-3 proteins are small, dimeric, acidic proteins that play a role in cellular signalling, apoptosis and cell cycle regulation (32-35). There are seven mammalian isoforms of 14-3-3 proteins (reviewed in (36)), which share a similar structure, consisting of nine α-helices that form the monomer, with the N-terminal domain being essential for the formation of the cup-shaped dimer (37,38). 14-3-3 proteins exist as both homo- or hetero-dimers and dimerization is essential for the ability of 14-3-3 proteins to form a complex with ligands and regulate ligand function (39-41). 14-3-3 dimers bind to their phosphorylated ligands via mode I (RSXpS/TXP) or mode II (RXXXpSXP) consensus sequences (42,43). 14-3-3 dimers bind to non-phosphorylated ligands via a mode III consensus sequence (44). Binding of 14-3-3 proteins to their ligands affects their localization, conformation, stability and function (33,45-49). 14-3-3γ and 14-3-3ε co-purify with the centrosomal fraction, (50,51) and loss of 14-3-3γ and 14-3-3ε leads to centrosome amplification in multiple cell lines due to the premature activation of CDC25C during interphase, leading to premature CDK1 activation and the increased phosphorylation of NPM1 at T199, which results in centrosome amplification (24).

While most 14-3-3 proteins bind to phosphorylated ligands containing one of two consensus sequences (42,43), not all 14-3-3 proteins bind to a given ligand in cells (52). This has led to two questions; how 14-3-3 proteins form complexes with their cognate ligands and how complex dissociation modulates ligand function. Therefore, in the present study, we focused on two conserved acidic amino acid residues in the peptide-binding pocket in 14-3-3 proteins, D129, and E136 in 14-3-3γ, to determine their contribution to ligand binding and dissociation. We find that while the expression of WT 14-3-3γ does not affect centrosome number, expression of the D129A mutant inhibits centrosome duplication. In contrast, expression of the E136A mutant leads to centrosome amplification. A double mutant, D129AE136A, shows a phenotype similar to WT 14-3-3γ. Further, we demonstrate that these effects are due to the differential binding of the 14-3-3γ mutants to NPM1. Modulation of the binding of 14-3-3γ and NPM1 can reverse the phenotypes observed upon expression of the 14-3-3γ mutants. Our results provide further insights into how 14-3-3 ligand complexes form and identify a novel mechanism by which 14-3-3γ regulates centrosome duplication.

## Results

### Acidic residues in the 14-3-3γ phospho-peptide binding pocket are required for accurate centrosome duplication

Most 14-3-3 ligands contain a mode I or mode II consensus motif that comprises of a phosphorylated Serine/Threonine residue (42,43). The overall structure of the protein, the binding pocket, and phospho-peptide interacting residues are very conserved in different 14-3-3 isoforms (53). A network of positively charged residues, K49, K120, R56, R127, and Y128 in 14-3-3ζ (K50, R57, R132 and R133 in 14-3-3γ) (figure 1A-C), anchors the phosphate ion in the peptide and these residues are conserved across isoforms. Charge neutralization or charge reversal of K49, R56, R127, and K120 result in partial or complete loss of interaction (54,55). Besides this strong positively charged cluster, there are backbone interactions mediated by N173 and N224 in 14-3-3ζ (N178 and N229 in 14-3-3γ) (figure 1A-C). Two hydrophobic residues, L172 and V176, line the pocket and pack against the hydrophobic residues of the amphipathic phospho-peptide. E180 located at the tail of helix 7 is hydrogen-bonded to the peptide main chain in mode I sequence. E180 also interacts with mode I Arginine in the 14-3-3ζ– AANAT complex (figure 1D) (47). Mutation of this conserved residue affects the binding of some 14-3-3 isoforms to Numb, while the interaction of other isoforms to the same ligand is unaffected (56).

**Figure 1.**
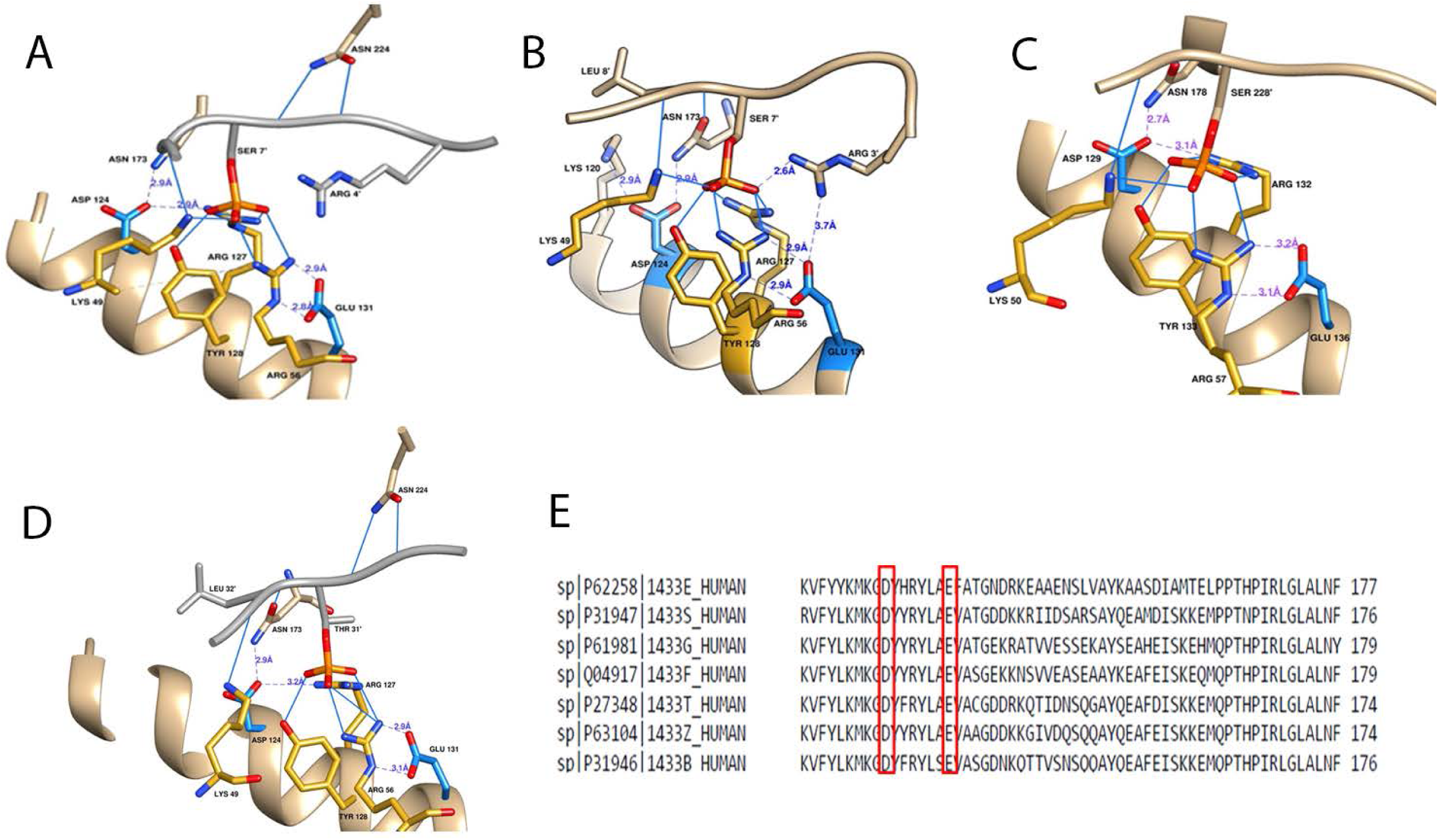
Sequence and structure conservation of phospho-peptide-14-3-3 protein interactions among the 14-3-3 isoforms. **(A-B)** Ribbon diagrams showing the presence of residues within 4A° of the two invariant residues Asp 124 and Glu 131 in the 14-3-3ζ mode I complex (A) or the 14-3-3ζ mode II complex (B). **(C)** Residues within 4A° of the two invariant residues Asp 129 and Glu 136 in 14-3-3γ phospho-peptide interaction. Note the absolute conservation of the structure of the network of positively charged residues that interact with the phospho-peptide in all three structures. **(D)** Ribbon diagram showing the presence of residues within 4A° of the two invariant residues Asp 124 and Glu 131 in the 14-3-3ζ – AANAT complex. **(E)** A sequence alignment showing the conservation of the Asp129 and Glu136 residues across different 14-3-3 isoforms.

There are other conserved residues in the binding pocket (figure 1E) and their role in ligand binding is unclear. Two conserved negatively charged residues attracted our attention - D124/129 and E131/136 in 14-3-3ζ or 14-3-3γ respectively. These residues lie within 4A°of the phospho-peptide in the crystal structures of 14-3-3ζ in complex with the mode I peptide (figure 1A-C). An examination of the binding pocket from the position of D129 or E136 elicits the striking observation that the positively charged residues that anchor the phosphate ion in the peptide are engaged in a network of interaction with these two residues (figure 1A-C). D129 anchors K49, K120, R127, and Y128. Similarly, E136 anchors R56, R127, and Y126. Besides, E136 is engaged in a weak salt bridge interaction with Arginine at the minus4 position in the mode II peptide and Arginine at the minus3 position in the mode I containing protein ligand (figure 1B and 1D), thereby suggesting that these conserved residues play an important role in ligand binding and might affect ligand function (43,47,56-58). These observations from the 14-3-3ζ crystal structures should be applicable to all 14-3-3 isoforms, including 14-3-3γ, because of the high degree of conservation in this region (figure 1E). The energetic contribution of these conserved residues to ligand binding has yet to be determined.

Given these observations, we decided to test if D129 and E136 in 14-3-3γ are essential for forming a complex with protein ligands and affecting ligand function. Previous work has demonstrated that 14-3-3γ is found in centrosome fractions (50) and that, loss of 14-3-3γ leads to an increase in centrosome number in human cell lines (24). We wanted to test if mutants of these conserved residues, D129 and E136 in 14-3-3γ, would have any effect on centrosome number. To that end, mOrange tagged WT 14-3-3γ or 14-3-3γ mutants were transfected into HCT116 cells and the 14-3-3γ constructs were expressed at equivalent levels (figure 2A). As shown in figure 2B-C, the expression of either mOrange or WT 14-3-3γ did not lead to a change in centrosome or centriole number with most mitotic cells showing two centrosomes, marked by Pericentrin, each containing 2 centrioles marked by Centrin2 (No. of puncta of Pericentrin:Centrin; 2:4). In contrast, a significant percentage of mitotic cells expressing the D129A mutant contained only one centrosome with two centrioles (19%) (No. of puncta of Pericentrin:Centrin2; 1:2), while a significant percentage of mitotic cells expressing the E136A mutant contained more than two centrosomes each with 2 centrioles (21%) (No. of puncta of Pericentrin:Centrin2; >2:4) (figure 2B-C). In contrast, mutation of the K50 residue to Alanine (K50A) did not affect centrosome number (figure S1A), though the K50E mutant has been reported to abolish ligand binding (40,59). A double mutant, D129AE136A, showed a phenotype similar to WT 14-3-3γ, an example of intragenic complementation (figure 2B-C). Similar results were observed in HaCaT and HEK293 cells (figure S1 B-C) suggesting that the phenotypes observed were independent of the cell line in which these experiments were performed. As each Pericentrin spot corresponded to two Centrin2 puncta, we used staining for Pericentrin to track centrosome number in all future experiments.

**Figure 2.**
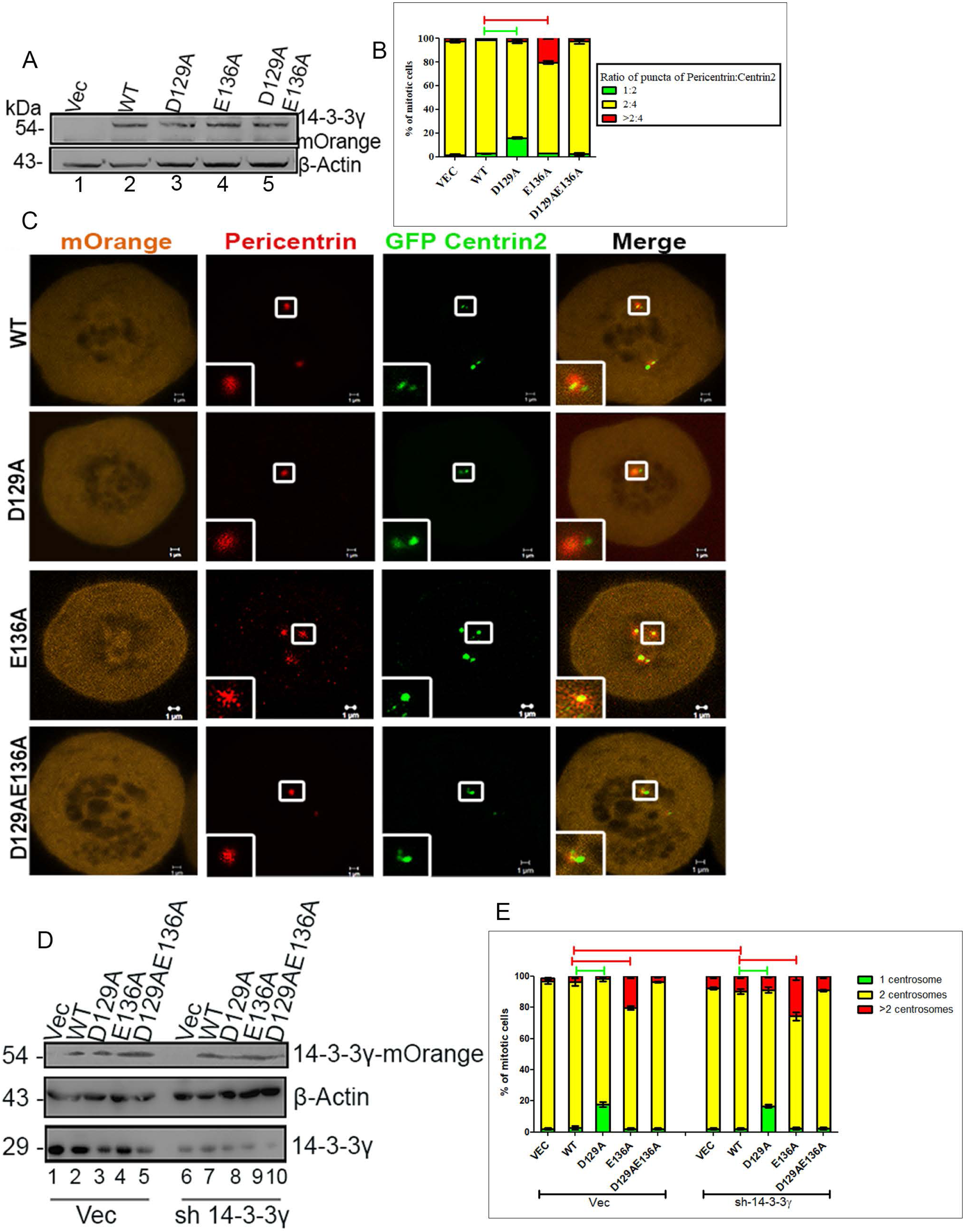
Effect of the 14-3-3γ mutants on centrosome number. **(A-C)** HCT116 cells were co-transfected with GFPCentrin2 and constructs expressing either the vector control (mOrange) or the mOrange tagged WT and mutant 14-3-3γ constructs. Protein extracts prepared from the transfected cells were resolved on SDS-PAGE gels followed by Western blotting with the indicated antibodies. **(A)** Note that all the 14-3-3γ proteins were expressed at equivalent levels (compare lanes 2-5). Western blots for β-actin served as a loading control. The transfected cells were fixed, permeabilized and stained with antibodies to Pericentrin (red) and counterstained with DAPI. The graph shows the percentage of mitotic cells with 1, 2 or >2 centrosomes and the number of centriole dots in each pericentrin dot **(B).** The panels in **(C)** show representative images and the inset is a magnified image of the area in the box. Note that the expression of D129A leads to an increase in the number of cells with a single centrosome and that the expression of E136A leads to centrosome amplification. Note that each pericentrin dot contains 2 Centrin2 dots. **(D-E).** The HCT116 derived vector control or 14-3-3γ knockdown clones (sh14-3-3γ) were transfected with the indicated constructs and centrosome number determined in mitotic cells. The graph shows the percentage of mitotic cells with 1, 2 or >2 centrosomes. All proteins were expressed at equivalent levels **(D)**. Note that D129A inhibits centrosome duplication in both the vector control and 14-3-3γ knockdown cells while E136A promotes centrosome duplication in both cell types **(E)**. In all the experiments the mean and standard error from at least three independent experiments were plotted and error bars denote standard error of mean. The horizontal bars indicate differences that are statistically significant (p<0.001). p-values are obtained using Student’s t-test. Original magnification 630X with 4X optical zoom. Scale bar indicates 2 µm unless mentioned. Molecular weight markers in kDa are indicated.

To test if the effect of the mutants is modulated by the presence of the endogenous protein, we wished to determine the effect of these mutants on centrosome number in the absence of 14-3-3γ as cells with a decrease in 14-3-3γ expression show centrosome amplification (24). We transfected the HCT116 derived vector control or 14-3-3γ knockdown cells with the constructs described above and determined centrosome number in mitotic cells. The 14-3-3γ knockdown cells showed an increase in the percentage of cells with multiple centrosomes (9%) as compared to the vector control cells as previously reported (p=0.0129) (figure 2D) (24). Expression of the D129A mutant in both the vector control and the knockdown cells resulted in a significant increase in the percentage of cells with a single centrosome (18% and 17% respectively) as compared to cells transfected with the WT construct or the vector control (figure 2D). In contrast, expression of E136A led to a significant increase in the percentage of cells with >two centrosomes in both the vector control and 14-3-3γ knockdown cells (20% and 25% respectively), while expression of the double mutant did not result in a significant change in centrosome number (figure 2D). All proteins were expressed at equivalent levels (figure 2E). These results suggest that the D129 and E136 residues play a role in regulating the contribution of 14-3-3γ to the centrosome cycle and that these mutants are dominant over the endogenous protein in human cells.

### The single centrosome phenotype is due to a defect in centriole biogenesis

There are three possibilities that can explain the presence of a single centrosome in cells expressing the D129A mutant. A defect in disengagement in G1 phase resulting in the mother centriole and daughter centriole remaining tethered to one another, a defect in centriole duplication during S phase or a defect in centrosome separation prior to the initiation of mitosis (figure 3A). The third possibility would suggest that 4 centriole dots would be observed in a single Pericentrin dot, which can be ruled out based on our previous results (figure 2B-C).

**Figure 3.**
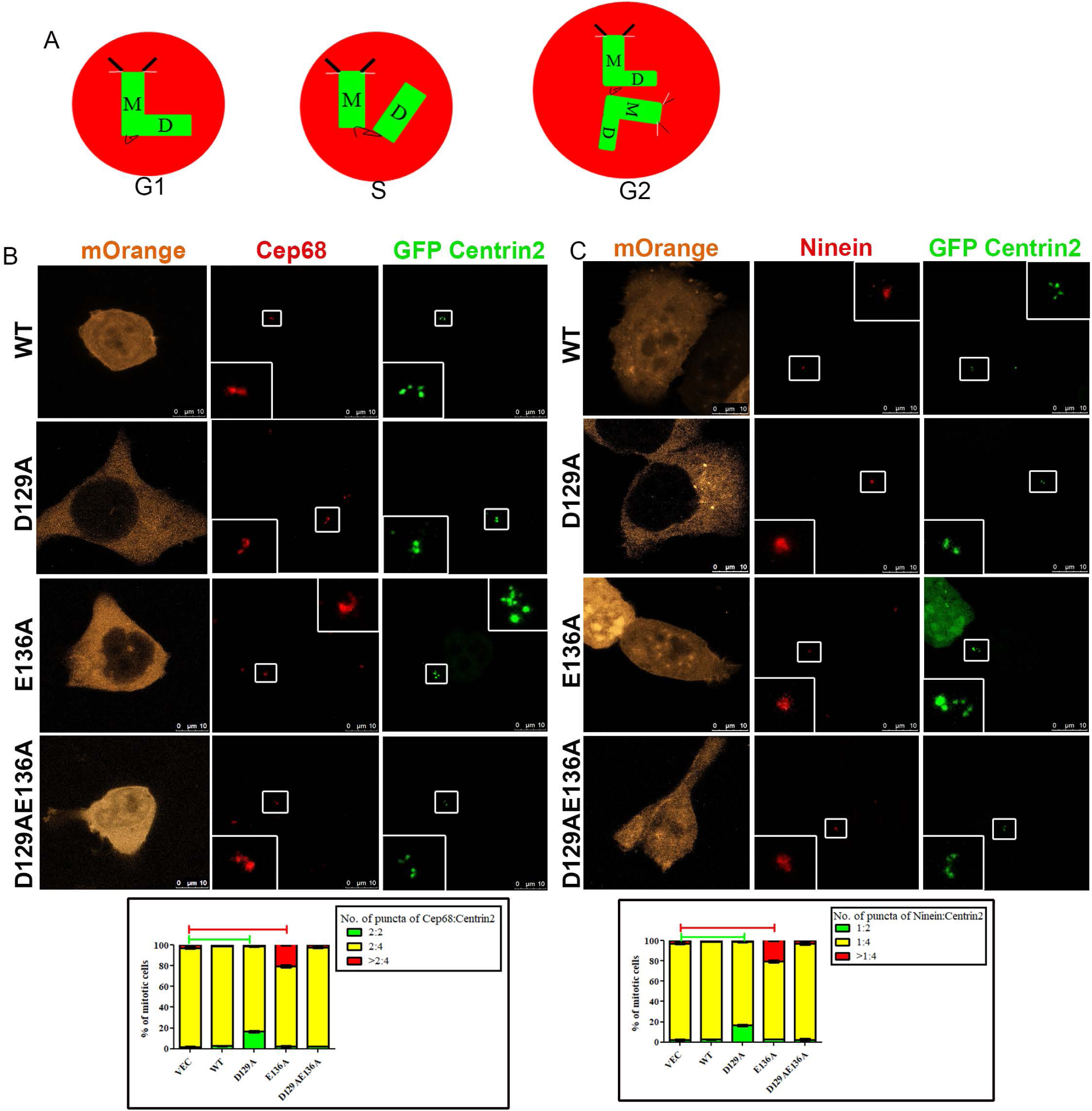
Centrosome organization in cells expressing the 14-3-3γ mutants. **(A)**. Cartoon showing the different causes for the presence of a single centrosome. **(B-C)** HCT116 cells were transfected with the indicated 14-3-3γ constructs and GFP Centrin2, synchronized in G2 with a CDK1 inhibitor, fixed and stained with Cep68 **(B)** and Ninein **(C)** and counterstained with DAPI. Representative images are shown and the insert is a magnification of the area in the box. **(B-C)**. Each single centrosome is associated with one Ninein dot and two Cep68 dots suggesting that disengagement has occurred but centriole synthesis has been inhibited. Original magnification 630X with 4X optical zoom. Scale bar indicates 10 µm unless mentioned. In all the experiments the mean and standard error from at least three independent experiments were plotted and error bars denote standard error of mean. The horizontal bars indicate differences that are statistically significant (p<0.001). p-values are obtained using Student’s t-test.

Centrosomes are licensed for duplication in G1 after disengagement of the mother and daughter centrioles, resulting in the formation of a G1-G2 tether, which consists of proteins such as C-Nap1, Rootletin and Cep68 (60-62) that loosely connects the two centrioles (reviewed in (7)). Therefore, if disengagement has occurred, each centriole will be associated with a pool of Cep68 in a single centrosome prior to centrosome duplication (No. of puncta of Cep68:Centrin2; 2:2). Post duplication, Cep68 will be present at each mother centriole (No. of puncta of Pericentrin:Centrin2; 2:4) until the beginning of mitosis when Cep68 degradation leads to centrosome separation (63). Therefore, if a centrosome with two centrioles in G2 has two Cep68 dots, it suggests that disengagement occurred without a subsequent initiation of centriole duplication. To test this hypothesis, HCT116 cells were co-transfected with the 14-3-3γ constructs and GFP Centrin2, arrested in G2 phase by treatment with the CDK1 inhibitor RO3306 and stained with antibodies to Cep68 (64). In cells expressing WT, E136A and the D129AE136A double mutant, each centrin pair was associated with one Cep68 dot (figure 3B). In contrast, cells expressing D129A that contained a single centrosome had one Cep68 dot associated with each centriole (figure 3B). A quantitative analysis of 100 transfected cells demonstrated that all the cells with a single centrosome containing 2 centrioles had a Cep68 dot associated with each centriole (figure 3B). These results suggested that in cells expressing the D129A mutant with a single centrosome, disengagement has occurred but centriole duplication is inhibited (No. of puncta of Cep68:Centrin2; 2:2).

To further characterize the single centrosome phenotype, we wished to determine whether the two centrioles were a mother and daughter pair. To test this, cells co-transfected with GFP-Centrin2 and the 14-3-3γ constructs were stained with antibodies to the mature mother centriole marker Ninein which localizes to the sub-distal appendages (65-67) post synchronization in G2 using RO3306, as Ninein is degraded during mitosis (68),. In cells expressing WT 14-3-3γ or the E136A and D129AE136A mutants, only one of the centriole pairs is associated with a Ninein dot (figure 3C). In contrast, in cells expressing D129A containing a single centrosome, the two centrioles are associated with one Ninein dot suggesting that the centriolar pair has been unable to divide. A quantitative analysis of 100 transfected cells demonstrated that the results were statistically significant (figure 3C). Taken together, all of the results reported above suggest that cells expressing the D129A mutant undergo disengagement but fail to initiate centriole duplication.

### 14-3-3γ inhibits centrosome licensing by regulating NPM1 function

Multiple kinases have been demonstrated to be required for the initiation of centriole duplication including Plk4 (27,28,69), CDK2 (70) and CDK1 (71,72). To determine if the expression of any of these kinases could result in a rescue of the defect in centriole duplication observed in cells expressing D129A, the 14-3-3γ constructs were co-transfected with myc-tagged Plk4, the WT and constitutively active forms of CDK2 and CDK1 into HCT116 cells. Centrosome number was determined in cells synchronized in mitosis by staining with antibodies to Pericentrin. As shown in figure S2A-F, expression of Plk4, CDK2 or CDK1 lead to centrosome amplification as demonstrated by the fact that over-expression of Plk4, CDK2 or CDK1 led to an increase in the percentage of cells with >2 centrosomes (figure S2A-F) as previously reported (12,21,24,72,73). However, the expression of these constructs did not stimulate centriole duplication in cells with a single centrosome, expressing the D129A mutant, as the percentage of cells with a single centrosome was unaltered in these cells. Thus, none of these kinases induced centriole duplication in cells with a single centrosome, expressing D129A.

Previous work from multiple laboratories has demonstrated that phosphorylation of a Threonine residue at position 199 (T199) in Nucleophosmin (NPM1) by either CDK1 or CDK2 is a licensing event for centriole duplication (22,74). Over-expression of either WT NPM1 or a phospho-mimetic mutant of NPM1 (T199D) can promote centrosome amplification while expression of a phospho-deficient mutant of NPM1 (T199A) inhibits centrosome amplification (24). To test if the expression of NPM1 could affect the phenotype observed upon expression of the 14-3-3γ mutants, the 14-3-3γ constructs were co-transfected with either the vector control or FLAG-tagged WT NPM1, T199A or T199D into HCT116 cells and centrosome number was determined as described. Expression of WT NPM1 or the T199D mutant resulted in a reduction in the number of cells with a single centrosome in cells expressing D129A (figure 4A-C). In contrast, the expression of T199A did not rescue the defect in centrosome duplication in cells expressing D129A but did inhibit the increase in centrosome number induced by expression of the E136A mutant (figure 4A-C). In all experiments, the NPM1 and 14-3-3γ proteins were expressed at equivalent levels (figure 4B). These results suggest that the D129A mutant might inhibit centriole duplication by preventing the release of NPM1 from the centrosome thus preventing centriole duplication.

**Figure 4.**
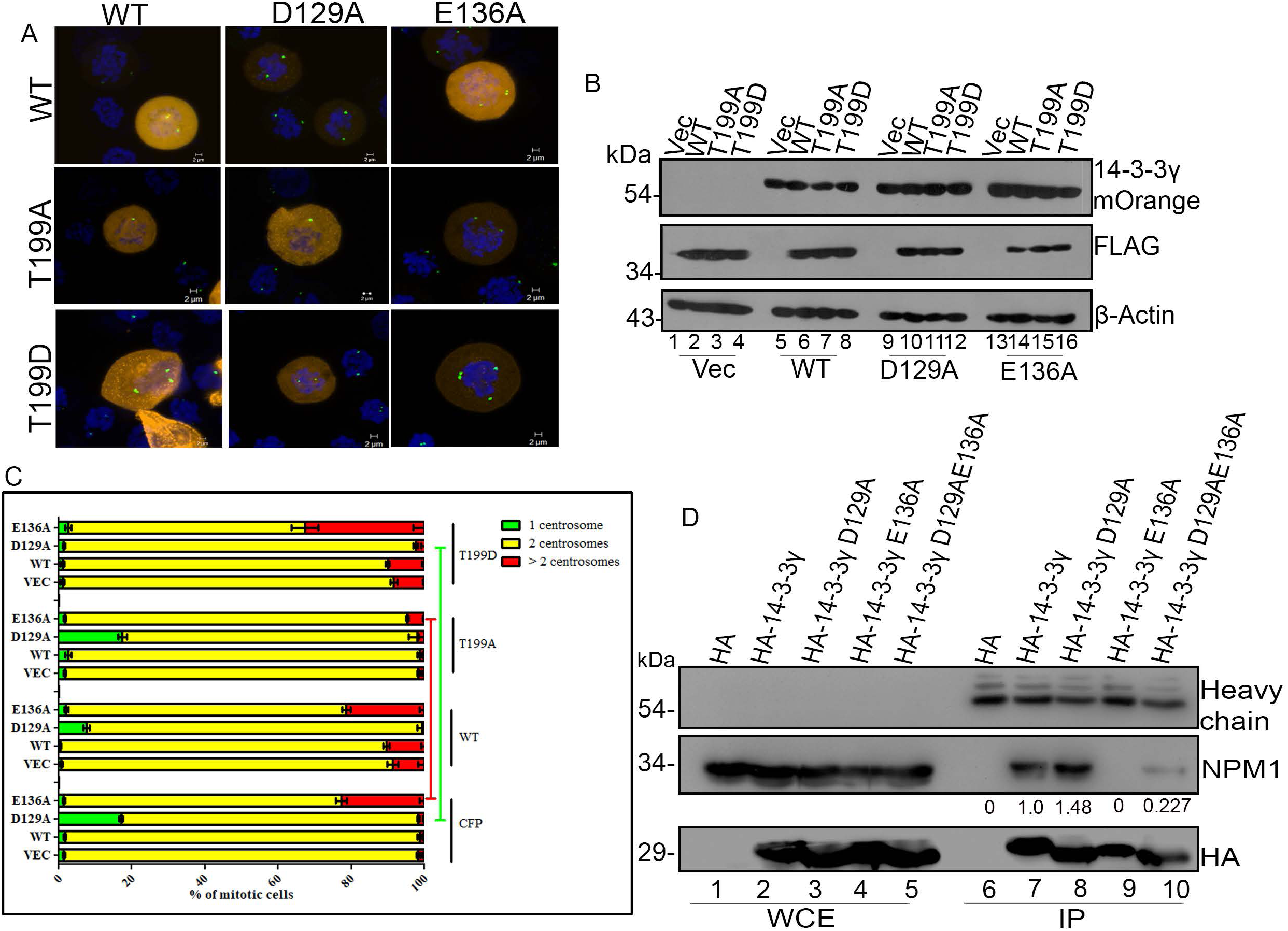
Expression of NPM1 results in an over-ride of the single centrosome phenotype observed in the D129A mutant. **(A-C)** HCT116 cells were transfected with the indicated 14-3-3γ and NPM1 constructs. Post-transfection the cells were arrested in mitosis and stained with antibodies to Pericentrin and counter-stained with DAPI. The representative images are shown **(A)** A Western blot analysis demonstrates that all proteins are present at equivalent levels (lanes 1-16) **(B). (C)** and the number of mitotic cells with 1 centrosome, 2 centrosomes or >2 centrosomes was plotted. Note that over-expression of NPM1 and T199D lead to a reversal of the single centrosome phenotype and over-expression of T199A leads to a reversal of the multiple centrosome phenotype. In all the experiments the mean and standard error from at least three independent experiments were plotted and error bars denote standard error of mean. The horizontal bars indicate statistically significant differences (p<0.001) and p-values are obtained using Student’s t-test. Original magnification 630X with 4X optical zoom. Scale bar indicates 2 µm unless mentioned. Molecular weight markers in kDa are indicated. **(D)** HCT116 cells transfected with the indicated constructs were lysed and subjected to immunoprecipitation with antibodies against the HA epitope. The reactions were resolved on SDS-PAGE gels followed by Western blotting with the indicated antibodies. Whole-cell extracts (WCE) were loaded in lanes 1-5 and the immunoprecipitation (IP) reactions in lanes 6-10. Note that while 14-3-3γ forms a complex with NPM1, E136A fails to form a complex with NPM1. D129A shows increased complex formation with NPM1 while D129AE136A forms a complex with NPM1 at reduced efficiency. The intensity of the co-immunoprecipitated NPM1 was determined and expressed as a fraction of that observed for WT14-3-3γ, which was set at 1. The numbers below the blot indicate the efficiency of co-immunoprecipitation for the different 14-3-3 constructs and NPM1.

NPM1 is phosphorylated by multiple kinases and functions as an oligomeric protein (22,75,76). To determine if phosphorylation at residues other than T199 could result in the same phenotype, other phospho-site mutants of NPM1 were transfected into HCT116 cells along with D129A. The other phospho-site mutants of NPM1 did not affect the percentage of cells with a single centrosome in cells expressing D129A (figure S3A). An oligomerization defective mutant of NPM1 (L18Q (77)) was also unable to decrease the percentage of cells with a single centrosome in D129A expressing cells, suggesting that the oligomerization of NPM1 was required for the ability of NPM1 to regulate centrosome licensing (figure S3B).

To determine if the different 14-3-3γ constructs could form a complex with NPM1, HA-epitope tagged versions of these proteins were transfected into HCT116 cells. 48 hours post-transfection, protein extracts prepared from these cells were incubated with antibodies to the HA epitope and protein G-Sepharose. The reactions were resolved on SDS-PAGE gels, followed by Western blotting with antibodies to HA or NPM1. As shown in figure 4D, the WT 14-3-3γ and the different 14-3-3γ mutants were expressed at equivalent levels in these cells. WT 14-3-3γ and D129A formed a complex with NPM1. However, E136A failed to form a complex with NPM1, suggesting that one reason why E136A expression induces centrosome over-duplication may be due to its inability to bind to NPM1. In contrast, the double mutant D129AE136A could form a complex with NPM1, though the degree of interaction is lower than that observed for WT or D129A (figure 4D).

Our results suggest that the D129A mutant binds to NPM1 with greater affinity than the WT protein. Further, the double mutant, D129AE136A is able to form a weak complex with NPM1 in contrast to E136A, which does not bind at all (figure 4D). In addition to a network of interaction involving the positively charged residues that bind the phosphate ion (figure 1), D124 (D129 in 14-3-3γ) also engages in interaction with a number of tyrosine residues located in and around 4A°on different helices. This implies that mutation of this residue would open up the protein facilitating the entry of the phosphorylated sequence. To test this hypothesis, the melting temperature of the WT and mutant 14-3-3γ proteins was determined (figure S3C). As compared to the WT 14-3-3γ, D129A is destabilized by 12°C while the E136A mutant was destabilized by 5°C (figure S3C). The D129AE136A double mutant was very unstable (figure S3C). Since the difference in fluorescence intensities between the folded and unfolded forms is minimal, the estimated Tm for the double mutant is not as reliable. Based on their stability to thermal denaturation, the proteins can be rank-ordered as WT>>E136A>>D129A>>> D129AE136A. The results suggest that a mutation that alters the D124/129 residue generates an open conformation of the protein facilitating easy access to the ligand.

Our results suggest that the phenotypes observed upon expression of the 14-3-3γ mutant might be due to the differential binding of these mutants to NPM1. Therefore, we decided to map the 14-3-3γ binding site in NPM1 and identified three Serine residues that could serve as potential 14-3-3 binding sites and altered them to Alanine (78). CFP tagged versions of these mutants or WT NPM1 were transfected into HCT116 cells. 48 hours post-transfection, protein extracts were prepared from these cells and incubated with bacterially expressed GST or GST tagged 14-3-3γ immobilized on Glutathione-Sepharose beads. The reactions were resolved on SDS-PAGE gels and Western blots performed with antibodies to GFP that also recognize CFP. WT NPM1 formed a complex with GST-14-3-3γ but not GST alone (figure 5A). Similarly, S218A and S293A formed a complex with GST-14-3-3γ, however, S48A failed to bind to GST-14-3-3γ, suggesting that S48 was required for the interaction of NPM1 with 14-3-3γ (figure 5A). None of the NPM1 constructs localized to the centrosome in our assays (figure S5).

**Figure 5.**
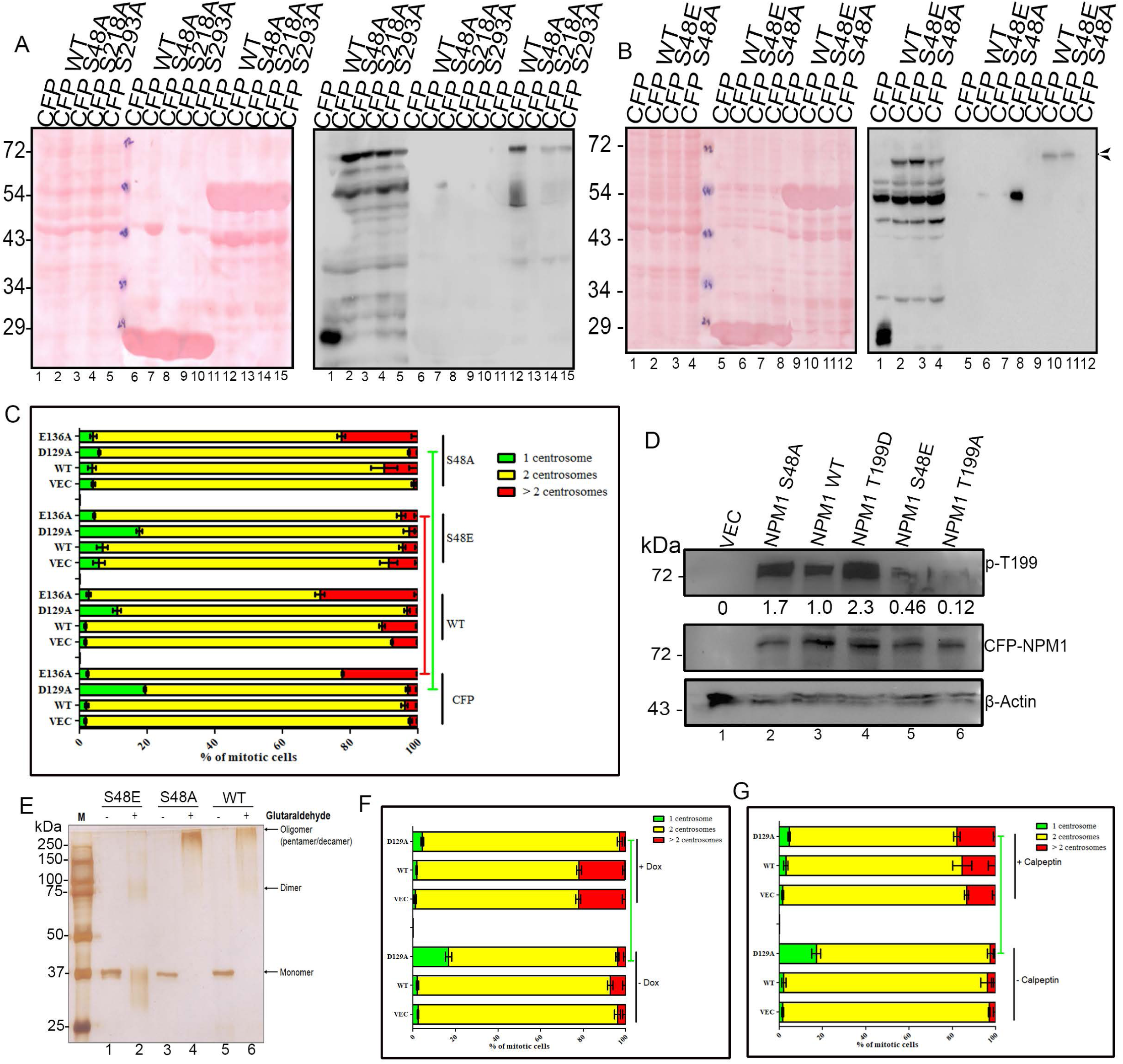
14-3-3γ inhibits NPM1 oligomerization and NPM1 function. **(A-B)**. HCT116 cells were transfected with the indicated NPM1 constructs. 48 hours post-transfection, protein extracts prepared from these cells were incubated with recombinant GST or GST-14-3-3γ immobilized on glutathione-Sepharose beads. The reactions were resolved on SDS-PAGE gels followed by Western blotting with the indicated antibodies. The panel on the left shows the Ponceau-S stain of the same membrane. In **(A)**, lanes 1-5 represent the WCE, lanes 6-10 represent pulldown with GST and lanes 11-15 represent pulldown with GST 14-3-3γ. In **(B)**, lanes 1-4 represent the WCE, lanes 5-8 represent pulldown with GST and lanes 9-12 represent pulldown with GST 14-3-3γ. Note that S48A fails to form a complex with 14-3-3γ, while S48E forms a complex with 14-3-3γ. **(C)** HCT116 cells were transfected with the indicated NPM1 and 14-3-3γ constructs. The cells were synchronized in mitosis with nocodazole, fixed, permeabilized and stained with antibodies to pericentrin and counterstained with DAPI. The percentage of mitotic cells with either one, two or more than two centrosomes is plotted. Note that expression of S48E inhibits the ability of E136A to promote centrosome hyper-duplication and that S48A reverses the single centrosome phenotype observed in cells expressing D129A. **(D)** HCT116 cells were transfected with the indicated NPM1 constructs. 48 hours post-transfection, protein extracts prepared from these cells were resolved on SDS-PAGE gels followed by Western blotting with the indicated antibodies. Western blots for β-actin serve as a loading control. Note that S48A shows increased phosphorylation on T199 while the phosphorylation is significantly reduced in the S48E mutant. The numbers indicate the intensity of the phospho T199 signal as compared to WTNPM1 which was set at 1. **(E)** Glutaraldehyde cross-linking assays were performed as described in materials and methods. Note that the S48E mutant cannot form higher-order oligomers. **(F)** HCT116 cells were transfected with the indicated 14-3-3γ constructs and a doxycycline-inducible constitutively active ROCK2 construct. The cells were cultured in the presence and absence of doxycycline and centrosome number determined in mitotic cells as described above. Note that the constitutively active ROCK2 construct only induces centrosome duplication in the presence of doxycycline. **(G)** HCT116 cells were transfected with the indicated 14-3-3γ constructs and treated with either the vector control or the ROCK2 activator Calpeptin and centrosome number determined in mitotic cells. Note that Calpeptin reverses the single centrosome phenotype in cells expressing D129A. In all the experiments the mean and standard error from at least three independent experiments were plotted and error bars denote standard error of mean. The horizontal bars indicate statistically significant differences (p=0.0001 **(C)** and p<0.03 **(F-G)**) and p-values are obtained using Student’s t-test. Molecular weight markers in kDa are indicated.

### 14-3-3γ inhibits NPM1 oligomerization and phosphorylation

Previous results have suggested that phosphorylation of S48 in NPM1 by Akt inhibits of the ability of NPM1 to form oligomers and that a phospho-mimetic mutant (S48E) failed to form oligomers *in vitro* (79). To determine if an S48E mutant of NPM1 could form a complex with 14-3-3γ, constructs expressing CFP fusions of WT, S48A, and S48E NPM1 were transfected into HCT116 cells followed by GST pulldown assays as described above. WT and S48E NPM1 formed a complex with GST-14-3-3γ but not GST alone, in contrast to S48A, which failed to form a complex with GST-14-3-3γ (figure 5B). Given these observations, we tested the ability of these mutants to affect centrosome number as previously described. All the NPM1 and 14-3-3γ mutants expressed at equivalent levels in these cells (figure S4A). Expression of S48A led to a decrease in the percentage of cells with a single centrosome and expressing D129A (figure 5C); while expression of S48E led to a decrease in the percentage of cells with multiple centrosomes upon expression of E136A (figure 5C). These results suggest that 14-3-3γ inhibits NPM1 function by inhibiting NPM1 oligomerization, a result consistent with our observation that an oligomerization defective mutant of NPM1 cannot reverse the single centrosome phenotype (figure S3B).

Given that phosphorylation of NPM1 at T199 stimulates the licensing of centrosome duplication (22,23), we determined the levels of T199 phosphorylation in the S48 mutants of NPM1 using a phospho-T199 specific antibody. The levels of phosphorylation of T199 on NPM1 were high in the case of the S48A mutant and reduced in the case of the S48E mutant protein (figure 5D). These results suggested that complex formation between NPM1 and 14-3-3γ inhibits the phosphorylation of NPM1 on T199 by CDK2 or CDK1. As the S48E mutation has been shown to inhibit NPM1 oligomerization (79), we performed oligomerization assays for WT and the S48 mutants of NPM1. All recombinant proteins were present at equivalent levels (figure S4B). The S48E mutant of NPM1 was defective for oligomerization as compared to the WT protein as previously reported (figure 5E) (79). The S48A mutant showed a small increase in oligomerization as compared to WT NPM1 (figure 5E). Similarly, histone chaperone assays demonstrated that the S48E had lower activity than WT NPM1 and the S48A mutant (figure S4C-D). These results suggest that complex formation between 14-3-3γ and NPM1 could potentially prevent NPM1 oligomerization and inhibit the phosphorylation of NPM1 at T199 resulting in a protein that cannot initiate the process of centrosome duplication.

### 14-3-3γ inhibits the NPM1-mediated activation of ROCK2, which is required for centrosome duplication

In addition to NPM1 phosphorylation at T199 being required for the initiation of centrosome licensing, it also seems to be required for the recruitment of the kinase ROCK2 to the mother centriole, which is required for the initiation of centriole duplication (80,81). Therefore, we wished to determine if activation of ROCK2 in cells expressing the D129A mutant would lead to a decrease in the number of cells with a single centrosome. To address this hypothesis, we transfected a doxycycline-inducible construct for a constitutively active ROCK2 kinase (ROCK2 CA) into cells with the different 14-3-3γ constructs and determined centrosome number in mitotic cells as previously described in the presence or absence of doxycycline (82). ROCK2 levels increased in the presence of doxycycline while the levels of the 14-3-3γ constructs did not change upon doxycycline addition (figure S4E). No change in centrosome number was observed in cells not treated with doxycycline (figure 5F). In cells treated with doxycycline, expression of ROCK2 CA led to a decrease in the number of cells with a single centrosome and expressing D129A, and an increase in centrosome amplification in cells transfected with the vector control or WT 14-3-3γ but not the D129A mutant (figure 5F). To confirm these results, cells transfected with the 14-3-3γ constructs were treated with the ROCK2 activator Calpeptin or the vehicle control and centrosome number determined as described (figure 5G). Treating cells with Calpeptin resulted in a decrease in the number of cells with a single centrosome in cells expressing D129A and an increase in centrosome amplification in cells transfected with the vector control or WT 14-3-3γ and the D129A mutant (figure 5G). Calpeptin treatment did not result in a change in ROCK2 levels but did lead to an increase in the phosphorylation of the ROCK2 substrate Myosin light chain IIC (figure S4F). Therefore, the inhibition of NPM1 function by 14-3-3γ inhibits ROCK2 activation at the centrosome, thus inhibiting centrosome duplication.

## Discussion

The results in this report identify a novel mechanism by which 14-3-3γ regulates centrosome duplication. 14-3-3γ forms a complex with NPM1, which prevents NPM1 oligomerization and the phosphorylation of T199 in NPM1 by CDK2. These events inhibit the ability of NPM1 to recruit ROCK2 to the centrosome, thus inhibiting centriole biogenesis and centrosome duplication. In addition, the results in this report identify the role of acidic amino acid residues in the 14-3-3 peptide-binding pocket in ligand association and potentially ligand release, thus further extending our understanding of how this important class of chaperones regulates cellular signalling pathways.

Based on the available crystal structures (figure 1 and figure S3A) and the thermal denaturation data (figure S3B), we have identified a possible role of the conserved acidic residues in 14-3-3 proteins, D124 (D129 in 14-3-3γ) and E131 (E136 in 14-3-3γ), in regulating complex formation with their ligands. While the crystal structure suggests that mutation of D129 may affect interaction of two key residues; N173 and R127 with the phospho-peptide, the experimental observations indicate other alternatives. The first is that the open D129A mutant is readily accessible to the protein ligand, a probable rate-limiting step in the interaction between 14-3-3γ and its ligands. The second is that the binding pocket in the D129A mutant is regenerated by an induced fit mechanism due to concurrent folding and binding of D129A to the ligand. The third is that in the absence of a potentially unfavorable D129 residue near the phosphate ion in the peptide, the network of positively charged residues is probably engaged in more stable interactions with the phosphate ion resulting in tighter binding. These results imply that D129 in 14-3-3γ is an important residue required for disengagement of the ligands from 14-3-3, which prevents dissociation of the complex thus affecting ligand function. Both the co-immunoprecipitation experiments (figure 4) and the consequences of the expression of these proteins on centrosome duplication (figures 2-3) are consistent with this hypothesis.

Analyses of the crystal structures of the mode I and mode II peptide bound 14-3-3 proteins (figure 1) and the crystal structure of NPM1 around the 14-3-3 binding site S48 (figure S3A), suggested that the 14-3-3 binding site in NPM1, which is a mode I peptide, is actually structurally similar to the mode II peptide. Based on this we have identified a potential role for the E136 residue in complex formation with NPM1. The Arginine in the minus three position in the NPM1 14-3-3 binding site might form a salt bridge with E136 in 14-3-3γ. The NPM1 structure suggests that the mode I motif would present itself in such a way that the Leucine at position 44 faces away from the amphipathic groove and the Arginine at position 45 (minus3 in the 14-3-3 mode I site) would tilt back into the binding pocket and form a salt bridge with E136 (figure 1). Mutation of E136 to Alanine might adversely affect protein binding, both due to this direct loss of contact and perhaps, due to the lack of appropriate orientation of the R56 in 14-3-3 in the peptide-binding pocket. This is consistent with the observation that the E136A mutant does not bind to NPM1 and induces centrosome over-duplication (figures 2 and 4). The thermal stability of E136A is similar to that of the WT 14-3-3γ indicating that it has a less important role in structural integrity as compared to that of D129A mutation. While the data support this conjecture, they do not explain why other residues do not compensate for the loss of E136. It is possible that E136 is strategically located at the wider end of the peptide-binding groove in 14-3-3γ and is the point of contact for protein entry.

In the D129AE136A double mutant, a far more open structure with an unstable fold and the absence of the key anchoring residue E136, should have resulted in increased dissociation of the complex and no net binding. However, this mutant formed a detectable complex with NPM1 and what is more, it resembled WT 14-3-3γ in its ability to regulate centrosome duplication suggesting intragenic complementation. The ability of D129A to reorient the residues at the binding pocket may enable the double mutant to form a complex with NPM1, although less efficiently than that of the WT protein. The faster dissociation would overcome the inhibitory tight binding seen with D129A mutant and restore centrosome duplication.

We propose the following model to explain how 14-3-3γ might form a complex with and dissociate from NPM1. The top panel in figure 6A shows the ligand-bound form of the 14-3-3γ WT and mutant proteins while the bottom panel shows the unbound form of the WT and mutant proteins. Note that the amphipathic loop is open in the case of the D129A and E136A mutants as indicated by the Tm (figure S3A). Also, note that in the more open D129A mutant, in the absence of a potentially repulsive negative charge, ligand-induced changes draw the positively charged cluster towards the negatively charged phosphate ion. This action ensures increased complex formation, which is probably due to an increase in the on-rate and a decrease in the off rate. In the E136A mutant, the protein fails to make the initial contacts, which prevents stable complex formation leading to an increase in the off rate.

**Figure 6.**
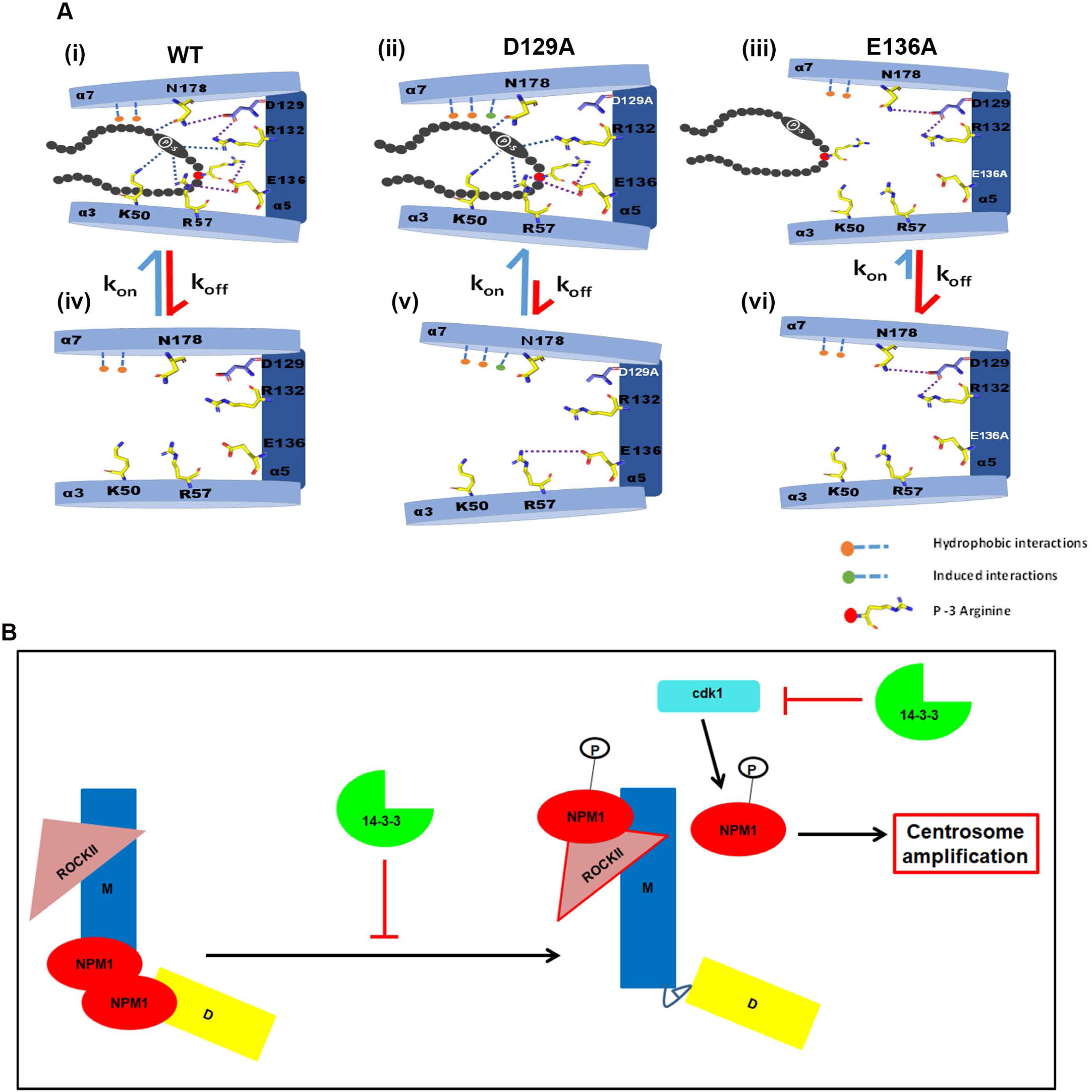
Model to explain how 14-3-3γ regulates NPM1 function and centrosome duplication. **(A).** Cartoon explaining the contribution of the acidic residues in the 14-3-3 peptide-binding groove to complex formation. **(B).** Cartoon explaining the role of 14-3-3 proteins in regulating different parts of the centrosome cycle.

Our results indicate that 14-3-3γ forms a complex with NPM1 and inhibits centrosome-licensing events initiated when CDK2 phosphorylates T199 in NPM1. Previously published work has suggested that NPM1 phosphorylation by CDK1 or CDK2 on T199 is required for the dissociation of NPM1 from the centrosome and the induction of centriole biogenesis (22-24,74). In addition, one function of NPM1 phosphorylated at T199 is the recruitment of activated ROCK2 to the mother centrosome, which is required for centrosome maturation and centriole duplication (80,81). Consistent with these observations, we have demonstrated that the single centrosome phenotype observed upon the expression of the D129A mutant can be over-ridden by expression of either a phospho-mimetic mutant of NPM1, T199D (figure 4), or by expression of a constitutively active form of ROCK2 or a small molecule that induces ROCK2 activation (figure 5). Further, the inactive T199A mutant of NPM1 can inhibit the ability of the E136A mutant of 14-3-3γ to induce centrosome amplification (figure 4) again suggesting that 14-3-3γ regulates centrosome licensing by inhibiting NPM1 function.

An analysis of the interaction between NPM1 and 14-3-3γ demonstrates that the association between 14-3-3γ and NPM1 requires an intact Serine residue at position 48 in NPM1. S48 in NPM1 can be phosphorylated by Akt and a phospho-mimetic mutant, S48E, fails to form higher-order oligomers in contrast to WT NPM1 (figure 5), which might affect the ability of NPM1 to serve as a protein chaperone (79). Our results demonstrate that the S48E mutant of NPM1 has poor oligomerization potential and decreased histone chaperone activity in contrast to the wild type protein or the phospho-deficient S48A mutant. Consistent with these observations, the S48E mutant can inhibit the centrosome amplification induced by the E136A mutant and the S48A inhibits the single centrosome phenotype observed in cells expressing the D129A mutant. 14-3-3γ might inhibit NPM1 function by inhibiting the phosphorylation of NPM1 on T199 by CDK1 or CDK2 as T199 phosphorylation is considerably decreased on the S48E mutant of NPM1 (figure 5). Another point mutant of NPM1, L18Q (77), that inhibits NPM1 oligomer formation cannot rescue the single centrosome phenotype observed upon D129A expression. Therefore, 14-3-3γ inhibits NPM1 function possibly by inhibiting higher-order oligomer formation and by inhibiting the phosphorylation of NPM1 on T199 by CDK1 or CDK2.

Phosphorylation of NPM1 at T199 by CDK2 results in the dissociation of NPM1 from the daughter centriole thus stimulating centriole biogenesis (22). NPM1 phosphorylation at T199 thus results in a re-localization of NPM1 to the mother centriole where it recruits and activates the kinase ROCK2 stimulating centrosome duplication (80,81). Despite multiple efforts, using antibodies reported in the literature (80,81) or epitope-tagged NPM1 constructs (this report); we have been unable to detect any NPM1 signal at the centrosome (figure S5). However, it is clear from our data that NPM1 expression can stimulate centrosome duplication in multiple cell types (this report and (24)) and that NPM1 mutants that fail to form a complex with 14-3-3γ induce centrosome amplification that is dependent on the presence of activated ROCK2 kinase. Given the chaperone functions of NPM1 (79), it is possible that it is required for the localization of ROCK2 to the centrosome upon CDK2 activation leading to centriole biogenesis and centrosome maturation.

Based on the results reported herein we would like to propose the following model for how 14-3-3γ regulates centrosome duplication (figure 6B). 14-3-3γ binding to NPM1 inhibits both the ability of NPM1 to form higher-order oligomers and the phosphorylation of NPM1 on T199 by CDK2. In addition, 14-3-3γ prevents centrosome re-duplication in S-phase by inhibiting CDC25C function as previously reported (24) and is required for the localization of Centrin2 to the centrosome (83). These results suggest that 14-3-3γ plays multiple roles in regulating centrosome duplication by preventing licensing prior to S-phase through inhibition of NPM1 and preventing reduplication later in the cell cycle by inhibiting CDC25C function.

In summary, the data in this report demonstrate that complex formation between 14-3-3 proteins and their ligands is a dynamic process and that complex formation is regulated by the orientation of conserved acidic amino acid residues in the peptide-binding groove. Further, our results suggest that 14-3-3γ regulates multiple facets of centrosome duplication and might regulate centrosome duplication and biogenesis in human cells. These results are important as 14-3-3 ligand complexes have been proposed to serve as possible therapeutic targets in the treatment of diseases such as cancer (24) and an understanding of the molecular basis underlying the complex formation and dissociation could allow the design of small molecules that target these complexes.

### Experimental procedures

#### Plasmids and constructs

The pGEX4T1 14-3-3γ WT plasmid has been described previously (52). pcDNA3 Plk4 (Sak) wt (Nigg HR9) was a gift from Erich Nigg (Addgene plasmid #41165; http://n2t.net/addgene:41165: RRID:Addgene_41165). Cdk2-HA was a gift from Sander van den Heuvel (Addgene plasmid #1884; http://n2t.net/addgene:1884; RRID:Addgene_1884).

The CDK1 WT and CDK1 AF constructs have been described previously (24). pSLIK CA ROCK2 was a gift from Sanjay Kumar (Addgene plasmid #84649; http://n2t.net/addgene:84649; RRID:Addgene_84649). pEGFPCentrin2 (Nigg UK185) was a gift from Erich Nigg (Addgene plasmid #41147; http://n2t.net/addgene:41147; RRID:Addgene_41147). An shRNA resistant WT 14-3-3γ cDNA was amplified (primers listed in Table S1) and cloned into pCMV mOrange digested with EcoRI and XhoI (New England Biolabs). The following mutant versions of 14-3-3γ, D129A, E136A and D129AE136A were generated by site-directed mutagenesis as described (84). The mutants were also cloned downstream of the HA epitope tag in pCDNA3 HA 14-3-3γ (52) by replacing the WT construct using BamHI and XhoI (New England Biolabs). WT NPM1 was amplified from a FLAG-NPM1 construct described previously (24). The amplified sequence was cloned into pECFP-N1 digested with XhoI and Bam HI (New England Biolabs) and the NPM1 point mutants were generated using site-directed mutagenesis (84). A web-based prediction server was used to identify possible 14-3-3 binding sites (78). All primer sequences are listed in table S1.

#### Cell lines and transfection

HCT116, HaCat and HEK293 cells were cultured as described previously (85). All the cell lines were STR profiled. Polyethyleneimine (PEI) (Polysciences, Inc.) was used as a transfection reagent as per the manufacturer’s instructions. To perform the immunofluorescence and Western blot assays, cells seeded on a cover-slip in a 35 mm dish were co-transfected with 1µg of the mOrange 14-3-3γ constructs and 1µg of the EGFP-Centrin2 or FLAG NPM1 or ECFP NPM1 constructs. The cells were harvested 48 hours post-transfection and Nocodazole treatment. For the Co-Immunoprecipitation experiments, HCT116 cells seeded in a 100 mm dish were transfected with 5µg of the HA-14-3-3γ constructs and harvested 48 hours post-transfection. Two dishes were used per transfection. For the GST pulldown experiments, 15µg of the ECFP-NPM1 were transfected per 100 mm dish and two dishes were used per transfection.

#### Antibodies and Western blotting

Primary antibodies for 14-3-3γ (Abcam #ab137106; dilution 1:500), β-actin (Sigma #A5316; dilution 1:5000), GFP (Clontech #632375; dilution 1:15,000), NPM1 (Invitrogen #325200; dilution 1:5000), phospho-T199 NPM1 (Abcam #ab81551; dilution 1:2000), ROCK2 (Santacruz #sc-398519, dilution 1:200), phospho-MLC2 (S19) (Cell Signalling #3671, dilution 1:1000), FLAG (Sigma #F-1804, dilution 1:500), HA (12CA5 hybridoma supernatant, dilution 1:100), phospho-CDK1 (Y15) (Cell signalling #9111, dilution 1:500), Centrin2 (Santacruz # sc-27793-R, dilution 1:500) and CDK1 (Cell Signalling (POH1) #9116, dilution 1:1000) were used for Western blot experiments. The secondary goat anti-mouse HRP (Pierce) and goat anti-rabbit HRP (Pierce) antibodies were used at a dilution of 1:2500 for Western blots. The following antibodies were used for immunofluorescence analysis; Pericentrin (Abcam, #ab4448, dilution 1:1000), α-tubulin (Abcam, #ab7291, dilution 1:200), Ninein (Cloudclone #PAC657Hu01 dilution 1:200) and Cep68 (Abcam #ab91455, dilution 1:200). Secondary antibodies (conjugated with Alexafluor-568, Alexafluor-488 and Alexafluor-633 from Molecular probes, Invitrogen; dilution 1:100) were used for immunofluorescence studies. For immunoblotting, samples were prepared in Laemmli’s buffer and separated on a 10% SDS-PAGE gel, transferred to nitrocellulose membranes and probed with antibodies to the indicated proteins. Western blots were imaged on a Bio-Rad ChemiDoc and 3 independent experiments were used for quantitation of protein levels as described (86).

#### Cell Synchronization

For enrichment of a mitotic population, cells were treated with Nocodazole (10µM #M1404 Sigma-Aldrich) for 18 hours (87). To induce a G2 arrest, cells were treated with a CDK1 inhibitor, RO3306 (9µM #SML0569 Sigma-Aldrich) (64,88). To test the effect of ROCK2 activation using Calpeptin, transfected cells were first synchronized to G1/S using Mimosine (80µM Sigma Aldrich (#M0253)) for 20 hours (87). 6 hours post-release from Mimosine, when the cells were at S phase, cells were treated with Calpeptin (5µM Santacruz (#202516)) for 30 minutes. Calpeptin was washed off and Nocodazole was added to synchronize cells in mitosis (87). The synchronization was confirmed using flow cytometry as described (24).

#### Immunofluorescence

For immunofluorescence analysis, cells were seeded on 22 mm coverslips. Post-transfection and synchronization, cells were fixed with 4% paraformaldehyde and permeabilized with 3% Triton X-100. After washes, the primary antibody was diluted in 3% BSA and cells were incubated in the primary antibody for 2 hours in a humidity chamber. Cells were incubated in secondary antibody for 1 hour. DNA was labeled using DAPI and cells were mounted on glass slides using Vectashield (Vector Laboratories H-1000). To observe centriolar organization using GFP-Centrin2 and Pericentrin, in cells expressing the different mutants, the cells were imaged on a Carl Zeiss LSM 780 system. 0.75 μm thin sections of the entire cell were captured. Images are represented as a projection of the entire Z stack. Cells were imaged at 1000X with a 4X digital zoom. Images were processed using the LSM Image browser. For all other experiments, images were acquired on a Leica SP8 confocal microscope at a 630X magnification with a 4X digital zoom. 0.75 μm thin sections of the entire cell were captured. Images are represented as a projection of the entire Z stack. Images were processed on the Leica LASX software.

#### Assaying centrosome number

Cells transfected with the various constructs were synchronized in mitosis as described above. The number of centrosomes in 100 mitotic cells expressing the mOrange-tagged constructs was determined in three independent experiments. Where noted the number of GFPCentrin2, Cep68 or Ninein foci were determined in 3 independent experiments.

#### GST pulldown and co-immunoprecipitation assays

EBC extracts of HCT116 cells were incubated with the various GST fusion proteins as indicated. The complexes were resolved on a 10% SDS-PAGE gel and Western blotted with antibodies against the indicated proteins. For the co-immunoprecipitation assays, 100 µL of anti-HA antibody (12CA5 hybridoma supernatant) was used to precipitate the HA-tagged 14-3-3γ proteins.

#### Purification of 14-3-3γ WT and mutants

14-3-3γ and the mutants D129A, E136A and D129AE136A were cloned in pETYONG and transformed into *E. coli* BL21 *codon* (*+*). Overnight grown culture was diluted 1:100 in fresh LB broth and allowed to grow at 37°C at 180 rpm till the absorbance at 260nm reached 0.4. The cells were then incubated at 18°C so as to reduce the temperature of the growing culture and then induced with 100 µM IPTG at 18°C overnight with constant shaking at 180 rpm. The cells were collected by centrifugation and washed once with lysis buffer (50 mM Tris-HCl pH 8.0, 150 mM NaCl, 10% glycerol, 20 mM Imidazole), re-suspended in lysis buffer and lysed by sonication. The cell lysate was centrifuged at 18000 rpm for 45 min at 4°C and the supernatant was loaded onto 2 ml of Ni^2+^ NTA resin in a column pre-equilibrated with lysis buffer. The bound protein was eluted with 50 mM Tris-HCl pH 8.0, 150 mM NaCl, 200 mM Imidazole and 10% glycerol and eluted fractions were analysed on SDS-PAGE. Fractions containing His-14-3-3γ were pooled and dialyzed against buffer containing 50 mM Tris-HCl, pH 8.0, 400 mM NaCl, 10% glycerol. The histidine tag was removed using His tagged TEV protease during the dialysis step. TEV was removed by loading the protein mixture on Ni^2+^ NTA resin and collecting the eluent. The eluted protein was first concentrated using centricon (3 kDa cut off) and was isolated by gel filtration chromatography on a Superdex200 column and purity confirmed by silver staining.

#### Purification of recombinant NPM1

The different His-tagged NPM1 proteins were purified by Ni-NTA based affinity purification method as per protocol previously (89,90). Briefly, *E. coli* BL-21 (DE3) competent cells were transformed with the respective plasmids, cultured at 37°C, 180 rpm and induced with 0.5 mM IPTG at O.D._600_ = 0.3-0.5. Cultures were grown for an additional 3 hours, harvested by centrifugation and the bacterial pellets stored at -80°C. Cell pellets were re-suspended in 30 ml of homogenization buffer (20 mM Tris-Cl pH = 7.5, 10% v/v glycerol, 0.2 mM Na-EDTA pH = 8, 300 mM KCl, 0.1% Nonidet P-40, 20 mM Imidazole, 2 mM PMSF, 2 mM β-Mercaptoethanol) and lysed by sonication. The cytosolic fraction was collected after centrifugation at 12000 rpm, 4°C for 30 min and incubated with equilibrated Ni-NTA beads (Novagen) at 12 rpm, 4°C for 3 hours. Beads were collected by centrifugation and washed with wash buffer (20 mM Tris-Cl pH = 7.5, 10% v/v glycerol, 0.2 mM Na-EDTA pH = 8, 300 mM KCl, 0.1% Nonidet P-40, 40 mM Imidazole, 2 mM PMSF, 2 mM β-Mercaptoethanol). Washed beads were packed in column and protein was eluted in elution buffer (20 mM Tris-Cl pH = 7.5, 10% v/v glycerol, 0.2 mM Na-EDTA pH = 8, 100 mM KCl, 0.2% Nonidet P-40, 250 mM Imidazole, 2 mM PMSF, 2 mM β-Mercaptoethanol). Purity and yield of the protein were analysed by resolving the purified proteins on a 12% SDS-PAGE gel followed by silver staining. Accordingly, suitable fractions were pooled and dialyzed against BC-100 buffer (20 mM Tris-Cl pH = 7.5, 10% v/v glycerol, 0.4 mM Na-EDTA pH = 8, 100 mM KCl, 0.1% Nonidet P-40, 0.5 mM PMSF, 9.8 mM β-Mercaptoethanol) to remove imidazole. Concentrations of the proteins were determined by gel estimation taking BSA as standard. Proteins were aliquoted, flash-frozen in liquid nitrogen and stored at -80°C.

#### Supercoiling assay

The nucleosome assembly activity of wild type Nucleophosmin (NPM1 WT), the phospho-deficient mutant (NPM1 S48A) and phospho-mimic mutant (NPM1 S48E) was assessed by the *in vitro* histone transfer or supercoiling assay as described previously, with a few modifications (91). Briefly, 200 ng of supercoiled G5ML plasmid was relaxed using *Drosophila* Topoisomerase I (dTopoI) in 1X assembly buffer (2X buffer composition: 10 mM Tris–HCl pH 8.0, 1 mM EDTA, 200 mM NaCl, and 0.1 mg/ml BSA (Sigma)) for 40 min at 37°C. In a parallel reaction, the histone chaperone protein, NPM1, was incubated with 350 ng of rat liver core histones in 1X assembly buffer in a total volume of 20 µl, at 37°C for 40 min for histone binding to the histone chaperone. This was followed by combining the two mixtures and incubating at 37°C for 40 min whereby histones are deposited onto the relaxed template by the chaperone, generating one negative supercoil per nucleosome assembly. The reaction was terminated by incubating in Proteinase K stop buffer (200 mM NaCl, 20 mM Na-EDTA, 0.25 µg/µl Glycogen, 1% SDS, 0.125 µg/µl Proteinase K (Sigma)) at 37°C for 30 min. Mixtures were deproteinized and DNA was extracted by phenol-chloroform-isoamyl alcohol (25:24:1, pH 8) method. Extracted DNA was electrophoresed in a 1% agarose gel in 1X TBE (111.4 mM Tris, 103.1 mM boric acid, 2 mM EDTA) at 50 V for about 12 h. The supercoiled DNA recovered due to the histone chaperone activity of NPM1, was quantified using Image J software and expressed as a percentage of the total supercoiled DNA used.

#### Glutaraldehyde cross-linking assay

Oligomerization property of NPM1 and its mutants was assessed by the Glutaraldehyde crosslinking assay as described previously, with some modifications (92). Briefly, 1 µg of NPM1 WT and mutant proteins were mixed with 0.05% v/v of Glutaraldehyde (Sigma) in a total reaction volume of 10 µl made up by Buffer H (20 mM HEPES-NaOH pH 7.9, 50 mM NaCl, 0.5 mM EDTA, 10% Glycerol, 0.5 mM PMSF) and incubated at room temperature for 10-15 min. The reaction was stopped by adding SDS-gel loading buffer to a final concentration of 1X and boiling at 90°C for 10 min. After SDS-PAGE, the proteins were visualized by staining of the gel with silver nitrate.

## Acknowledgements

We thank Dr. Erich Nigg, for his kind gift of the EGFP C2 Centrin2 construct and the ACTREC flow, microscopy and digital imaging facilities for help with the experiments in this report. The anti-NPM1, anti-phospho-NPM1 (T199) antibody was a kind gift from Prof. Tapas K. Kundu, JNCASR, Bangalore. SD^3^ is supported by University Grants Commission (UGC), India and TKK is a Sir J. C. Bose Fellow. The work is funded by grants from the Department of Biotechnology, Govt. of India (BT/PR4181/BRB/10/1147/2012 and BT/PR8351/MED/30/995/2013) to SND.

## Conflict of interest

The authors declare that they have no conflicts of interest with the contents of this article.

